# Simulating groundcover community assembly in a frequently burned ecosystem using a simple neutral model

**DOI:** 10.1101/635169

**Authors:** E. Louise Loudermilk, Lee Dyer, Scott Pokswinski, Andrew T. Hudak, Benjamin Hornsby, Lora Richards, Jane Dell, Scott L. Goodrick, J. Kevin Hiers, Joseph J. O’Brien

**Affiliations:** USDA Forest Service, Southern Research Station, Center for Forest Disturbance Science, Athens, GA, USA; University of Nevada, Reno, Department of Biology, Reno, NV, USA; Tall Timbers Research Station and Conservancy, Tallahassee, FL, USA; USDA Forest Service, Rocky Mountain Research Station, Forestry Sciences Laboratory, Moscow, ID, USA

## Abstract

Fire is a global process that drives patterns of biodiversity. In frequently burned fire-dependent ecosystems, surface fire regimes allow for the coexistence of high plant diversity at fine-scales even where soils are uniform. The mechanisms on how fire impacts groundcover community dynamics are however, poorly understood. Because fire can act as a stochastic agent of mortality, we hypothesized that a neutral mechanism might be responsible for maintaining plant diversity. We used the demographic parameters of the Unified Neutral Theory of Biodiversity (UNTB) as a foundation to model groundcover species richness, using a southeastern U.S. pine woodland as an example. We followed the fate of over 7,000 individuals of 123 plant species for four years and two prescribed burns in frequently burned *Pinus palustris* sites in NW FL, USA. Using these empirical data and UNTB-based assumptions, we developed two parsimonious autonomous agent models, which were distinct by spatially explicit and implicit local recruitment processes. Using a parameter sensitivity test, we examined how empirical estimates, input species frequency distributions, and community size affected output species richness. We found that dispersal limitation was the most influential parameter, followed by mortality and birth, and that these parameters varied based on scale of the frequency distributions. Overall, these nominal parameters were useful for simulating fine-scale groundcover communities, although further empirical analysis of richness patterns, particularly related to fine-scale burn severity is needed. This modeling framework can be utilized to examine our premise that localized groundcover assemblages are neutral communities at high fire frequencies, as well as examine the extent to which niche-based dynamics determine community dynamics when fire frequency is altered and spatial fire intensity patterns differ.

## 1 Introduction

Wildland fire is a globally critical process for maintaining biodiversity in many terrestrial ecosystems (Pausas and Ribeiro 2017). Surface fire regimes are particularly important, where frequent low-intensity fires often maintain the highest levels of floral and faunal diversity (Bond and Keeley 2005, Mitchell et al. 2009, Pausas and Ribeiro 2017). These frequently burned ecosystems are a significant global biome (Staver et al. 2011), thus understanding how fire structures patterns of diversity is a critical need. Longleaf pine (*Pinus palustris* Mill.) woodlands of the southeastern USA are dependent on frequent fire with burns occurring as frequently as every 18 months. Another characteristic of these systems is the high plant species richness and endemism that occurs at fine scales (Hardin and White 1989, Barnett 1999, Dell et al. 2019) within the groundcover plant community. The relationship between high fire frequency and high groundcover diversity is well known, but the mechanisms driving this connection still remain elusive. Previous work (Platt et al. 1988, Kirkman et al. 2001, Pausas and Ribeiro 2017) has focused on documenting correlations between species diversity, fire return intervals, soil edaphic gradients (i.e. productivity), and seasons of burn. Other studies have tested the competitive effects of dominant guilds (bunchgrasses or shrubs) on rare species in these frequently burned systems and found no competitive effects (Myers and Harms 2009), while others found facilitation effects (Iacona et al. 2012). These studies do not however, account for microscale spatial demographics that might structure the groundcover community, and do not attempt to project richness patterns based on demographic parameters. Furthermore, many small plant species are found in high densities on very uniform sandy soils (up to 40-50 within 1 m^2^, Peet and Allard 1993, Kirkman et al. 2001). If niche-based processes were structuring these communities, resource-partitioning for dozens of species would occur within just a few meters across these resource-poor soils.

Research on ecological community assembly and biodiversity has been invigorated by the Unified Neutral Theory of Biodiversity (UNTB, Hubbell 2001, Chave 2004, Matthews and Whittaker 2014, Missa et al. 2016). The theory has stimulated modeling and empirical studies examining the relative roles of stochastic processes versus competitive exclusion and niche differentiation in driving patterns of species richness and community assembly, particularly among plants (McGill 2003, Volkov et al. 2003, Chave 2004, Wootton 2005, John et al. 2007, Rosindell et al. 2011). UNTB assumes that individuals of different species within a trophic level are functionally equivalent and posits that communities of these species are structured by random demographic processes, such as birth, death, and dispersal limitation, and at longer time scales, speciation and extinction.

For many ecosystems, both neutral processes and niche differentiation affect community structure (Gravel et al. 2006, Adler et al. 2007, Matthews and Whittaker 2014) with the UNTB treated primarily as a null model that captures the impact of unknown mechanisms (Alonso et al. 2006, Rosindell et al. 2012). We hypothesize that the high species richness of frequently burned ecosystems may be an actual consequence of these neutral processes at smaller temporal and spatial scales because fire plays two crucial roles. One, repetitive fire (1-5yr. intervals) suppresses competition of fast-growing woody plant species (Barnett 1999, McGuire et al. 2001) and acts as a competitive filter (Pausas and Verdú 2008, Myers and Harms 2011), and two, fire causes random mortality at fine-scales (Wiggers et al. 2013, O’Brien et al. *2016a*). These random mortality events are a defining process of the UNTB. Fire driven mortality in these systems occurs at the scale of an individual plant (Wiggers et al. 2013, O’Brien et al. *2016a)*, because the relevant scale of variability in fire intensity of these surface fires is also found at this same fine scale (Hiers et al. 2009, Loudermilk et al. 2012). These neutral dynamics however are usurped when fire is removed from these systems as predictable changes in the plant community occur, with faster growing fire sensitive species released and competition drives the shift to woody plant dominance (Brockway and Lewis 1997, Glitzenstein et al. 2003). Ultimately, the relative importance of mortality and other neutral parameters (birth, dispersal limitation) in these or similar groundcover communities associated with fire adapted forests remains unexplored.

Despite criticisms and attempts to reject the UNTB, the utility of estimating and examining variation in demographic parameters in the context of neutral contributions to biodiversity is clear (Rosindell et al. 2012). Whether or not it is treated as a null model, UNTB allows for deviations from neutrality, and different demographic parameters are more sensitive to change. Thus, it is important to examine how changes in neutral parameters affect diversity predictions across different communities and across different scales (Hubbell 2001, Matthews and Whittaker 2014). Properly quantifying variation in neutral parameters within and among modeled communities and at multiple scales requires careful development to ensure accurate predictions of species richness. There are a number of approaches for examining such variation, including sensitivity analyses, which may be used to quantify the relative differences in sensitivity between parameters (Hamby 1994, Saltelli et al. 1999). These analyses are used with newly developed models to provide insight into how parameters vary and to generate guidance on appropriate model interpretation and future directions.

In this study, we explored how neutral processes (Hubbell 2001) might be the mechanism maintaining high groundcover diversity when exposed to frequent fire. We explored UNTB-based parameters within a modeling environment using empirical observations. We utilized four years of plant census data with individual plants mapped at very fine-scales (100 cm^2^), following their fates before and after two low intensity prescribed burns in a northwest FL, USA longleaf pine ecosystem. We estimated UNTB-based parameters for two parsimonious autonomous agent models, which were distinct by spatially explicit and implicit local recruitment processes. We employed parameter sensitivity tests within our two models to examine how parameter estimates affected output species richness. We also examined how varying input species frequency distributions and community size assumptions of the model influenced parameter sensitivity across models.

## 2 Materials and Methods

### 2.1 Study Site

This study was conducted at Eglin Air Force Base (EAFB), FL from 2014 to 2016. EAFB, the former Choctawhatchee National Forest, is located in northwest Florida, USA (30° 34’23.50”N, 86° 34’48.64”W). The site includes nearly 180,000 ha of longleaf pine forests, which contains over half of the remaining old growth stands of this forest type (Varner et al. 2000, Holliday 2001). The study sites were within the Southern Pine Hills District of the Coastal Plain Physiographic Province with deep, well-drained sandy soils (Overing et al. 1995). The climate of the area is subtropical with mean annual temperatures of 19.7°C and mean annual precipitation of 1580 mm (Guo et al. 2016). Elevations were 52 to 85 m above sea level. Vegetation was dominated by a longleaf pine overstory, with a sparse midstory of various deciduous oaks, e.g., *Quercus laevis* Walter, *Q. margaretta* Ashe, *Q. incana* Bartram, *Q. germinata* Small.

### 2.2 Data Collection and Prescribed Burning

We established our plots within existing EAFB monitoring plots in two habitat types— sandhills and flatwoods. The sandhills habitat is xeric with deeper sands and at a higher elevation than the mesic flatwoods habitat. The flatwoods have more productive soils than the sandhills. In each habitat type, we established 15 random 1m × 3m plots for a total of 30 plots. Every Fall from 2012 to 2016, yearly records of all living plants within each plot (sandhills n=15 and flatwoods n=15) were mapped using a 10 cm × 10 cm grid coordinate system. All plant individuals were recorded by species. The resulting plant dataset includes four intervals of data comprising of 123 plant species (**Table S1)** and over 7000 individuals. Two experimental prescribed burns were conducted in Spring 2013 and 2015. The fires were conducted as part of normal forest management operations at EAFB (O’Brien et al. 2016a, O’Brien et al. 2016b).

### 2.3 Autonomous Agent Modelling

To model groundcover community dynamics, we developed one spatially explicit and one spatially implicit neutral autonomous agent simulation model, respectively the ‘spatial model’ and ‘non-spatial model,’ using the python programming language (v. 2.7.11 Python Software Foundation). As the input data were collected in frequently burned areas of EAFB with a two-year return interval, we assumed that fires were implicit in both models. This is realistic because after two years without fire, species richness begins to decline in these ecosystems (Brockway and Lewis 1997, Glitzenstein et al. 2003). A conceptual model of model inputs and processes are found in **Fig. 1.**

**Fig. 1.**
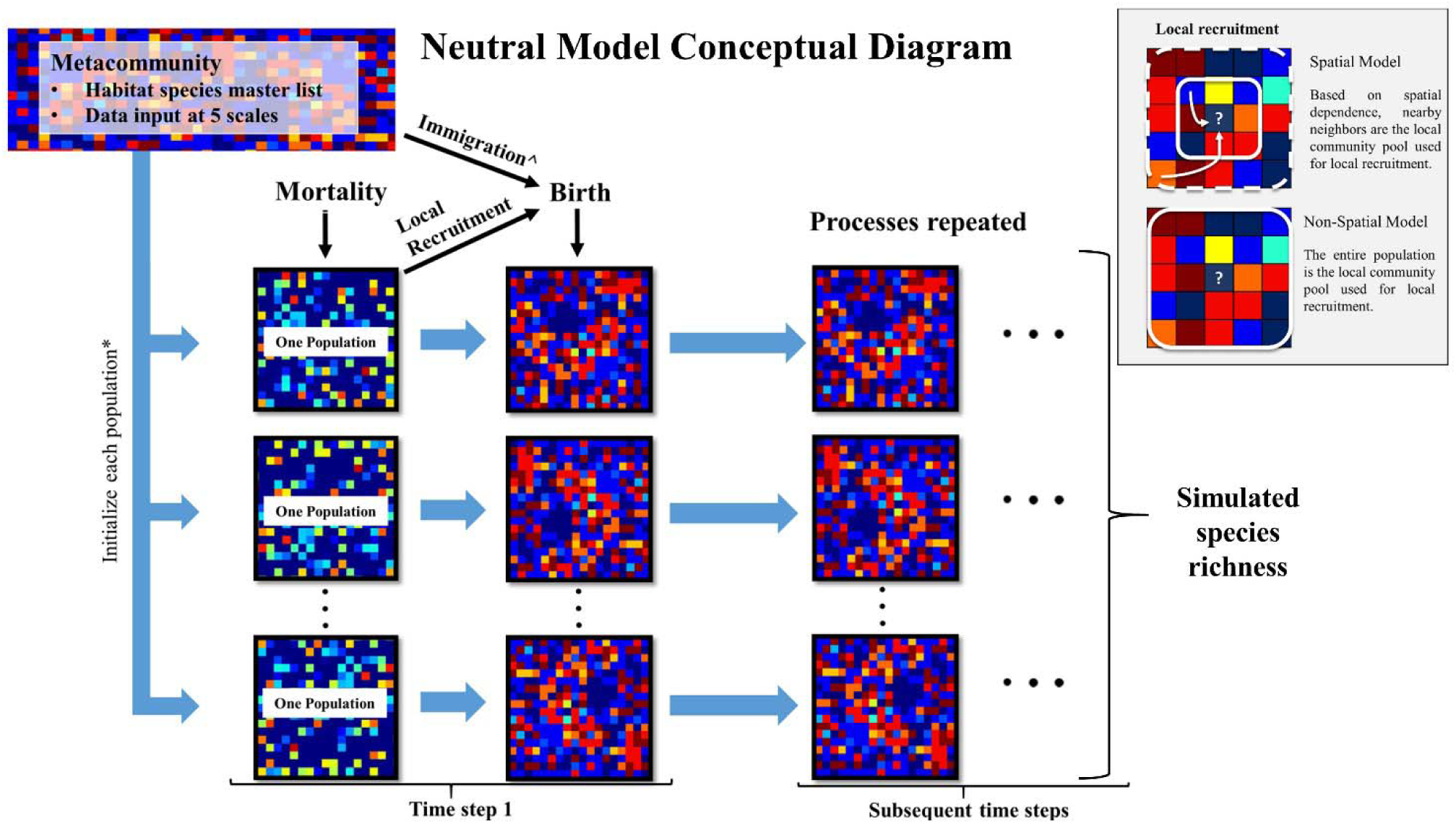
Conceptual diagram of this study’s neutral model used to simulate groundcover species richness in a frequently burned landscape. The metacommunity is the groundcover species frequency distribution within each longleaf pine habitat (flatwoods and sandhills) as recorded in this study at EAFB (Fig. S1, S2). These data were used as input at 5 scales (described in text) to initialize each population simulated in the model. Each individual experiences a probability of mortality. Then empty cells experience a probability of birth, where species are recruited from either the local community pool or metacommunity pool, the later through immigration. The box insert describes how local recruitment is simulated in the spatial and non-spatial versions of the model. The resulting populations are used as the starting population for the subsequent timesteps. *The size of each population and number of populations are determined by the community size parameters: area and number of areas; ^ If immigration occurs during a ‘birth’ event, the metacommunity pool is used in place of the local community (population) pool for recruitment.

UNTB requires a metacommunity species pool, a local community species pool, and community level birth, death, and immigration (dispersal limitation) rates. To estimate the distribution of plant species found across each habitat, we combined all plant data from the burned areas to create one input species distribution for each habitat (**Fig. S1**). These metacommunity species distributions (“the metacommunity”) were used to populate the initial distribution of plant species within each simulated area by randomly choosing species from this distribution. The metacommunity also provided a source for recruitment that occurred from immigration only (not the local community) throughout each simulation.

We simulated individual plants through a presence-absence approach, where each individual was subjected to a yearly mortality risk and empty cells to recruitment potential. The probability of birth (recruitment) and death were produced using a random number generator and a set seed with no density dependence effects on death. We used a lattice-base modeling approach with a 1 year time-step, where only one individual could occupy a cell in the lattice. Each cell was 100 cm^2^, the same scale at which individuals were sampled in the field. The spatial model was distinguished from the non-spatial model by the implementation of periodic boundaries, which eliminated edge effects on spatially dependent recruitment.

For the non-spatial model, local recruitment was a function of the probability of birth, where species were sampled from the frequency distributions for each simulated area (i.e. the local community pool). For the spatial model, local recruitment was a function of the probability of birth and the local neighborhood of individuals. Here, we used a negative exponential function within the NumPy (v1.11.3) module for python, to determine which individual from the neighboring community will recruit into the focal cell. We set a maximum distance of five cells in all directions from the focal cell for local dispersal to standardize the local recruitment pool. This sets a maximum dispersal distance to 0.71 m, the distance diagonal from the focal cell. This includes an area up to 1.1025 m^2^ (1.05 m × 1.05 m), which represents the local community pool at any given “birth” event. Although seed dispersal is quite variable among these groundcover species (Stamp and Lucas 1990), we found this seeding distance to be comparable to a similar longleaf pine reference site, where most species had restricted seed dispersal mechanisms (Kirkman et al. 2004).

### 2.4 Model Parameterization and Sensitivity Analysis

Mortality and birth rates were estimated from the plant monitoring data. We used plant data from the 2012 to 2014 and 2014 to 2016 surveys because it was a consistent 1.5 year time period between burns, including the 2012 survey, which initially burned in Spring 2011. An individual plant of an identified species was considered dead if it was not found in its original cell in the next census. Recruitment occurred when a plant was newly found in a cell in the next census. Sensitivity analyses were then utilized because the resulting mortality and birth rates across plots, years, and habitats were highly variable, and when these values were applied to the models had confounding effects on model outputs. We employed the Fourier Amplitude Sensitivity Test (FAST) as defined by (Saltelli et al. 1999) to provide an unbiased approach to assess the effects of each range of parameter values using a concise method. The FAST was applied in the python library SALib (v.1.0.2) to examine the first order effects of model parameters on simulated species richness. FAST quantifies the sensitivity of multiple parameters simultaneously by assigning each input parameter a frequency at which to oscillate across a range of input values. Decomposing the model outputs using these same frequencies allows for the extraction of the influence of each input parameter on the model results. This allows model sensitivity to be assessed using far fewer simulations than a complete exploration of the parameter space. Using this method, a unique set of input parameter values was created for each model run using the empirical data.

Five model parameters were used in the FAST; the three UNTB-based or neutral parameters (mortality, birth, and immigration), and two more exogenous-type parameters that influence the spatial boundaries in which the model runs, namely area (size of each population) and number of areas (or populations). The two additional parameters (i.e. community size parameters) were included because they were found to drive community size during model initialization, recruitment, and immigration, all influencing output species richness. Input parameter sets were created for each of these five model parameters within each habitat.

The values used as input parameter sets in the FAST include birth rate (0.2-0.6), mortality rate (0.2-0.6), immigration rate (0.01-0.25), area simulated (1m^2^ – 25m^2^), and number of areas simulated (1-30 areas). We chose a range of plot (area) sizes and numbers of plots (areas) used to estimate species richness in these small plant communities (Hardin and White 1989, Peet and Allard 1993, Kirkman et al. 2001, Dell et al. 2017) to explore their effects on simulated outputs. Our immigration rates are a range of values that represent a high migration system (>0.1) and low migration system (<0.1), which affects rates of species turnover (Volkov et al. 2003). In SALib, 100 samples for each of the five parameter sets were created. This resulted in 500 unique parameter combinations used to simulate each habitat and scale of input for the spatial and non-spatial model. To include variability, three replicates using three different random number seeds were simulated. Our target output variable was “normalized richness,” which is the proportion richness change from time-step one to the ending time-step of 50, when model surpassed any instability.

Furthermore, we explored issues associated with initial conditions to assess their impact on simulated species richness. This was performed because it has been found that initial conditions, particularly related to scale of the species frequency distributions, can influence simulations of neutrality (Hubbell 2001, Matthews and Whittaker 2014). As such, we examined effects of the scale of inputs by coarsening the initial species frequency distributions using a presence-absence approach across various resolutions, resulting in inputs at the 10 cm × 10 cm (original resolution), 20 cm × 20 cm, 50 cm × 50 cm, 100 cm × 100 cm, and 200 cm × 200 cm scales. Within each resulting coarser size cell, species presence was recorded. These species distributions (**Fig. S1**) were used as initial inputs to assess effects of scaled initial conditions on the FAST simulations.

In SALib, the simulation outputs were analyzed across the 500 parameter combinations and 20 scenarios (2 models × 2 habitats × 5 scales of inputs). We focused on the first order effects of the parameters, illustrating the percent of the variance in normalized richness which is explained by each parameter alone. This allowed us to focus on individual parameter sensitivity to a multi-faceted combination of initial conditions and model processes.

## 3 Results

Our autonomous agent models simulated groundcover species richness distributions across two longleaf pine habitats (**Fig. 2, S2**). The results revealed there was little difference in outputs between habitats (**Fig.s 2, 3, S2, S3**), which is a result of the similarity in size and shape of their input species frequency distributions or their metacommunities (**Fig.s S1**). For brevity, we focus on the sandhills habitat results here, but include all flatwoods outputs in **Fig.s S2 and S3**. The FAST illustrated that the direct first order effects of varying the mortality and birth parameters (from 0.2 to 0.6) contributed to less than 5% of the model variance (**Fig. 3**). Immigration was the most sensitive of the neutral parameters, varying from about 5% up to 20%. The community size parameters (number of areas and area) were similarly sensitive, varying also up to 20%, and illustrated the most sensitivity between replicates (**Fig. 3**). There were little differences between species richness distributions and parameter sensitivities between the spatial and non-spatial models (**Fig. 2, 3**).

**Fig. 2.**
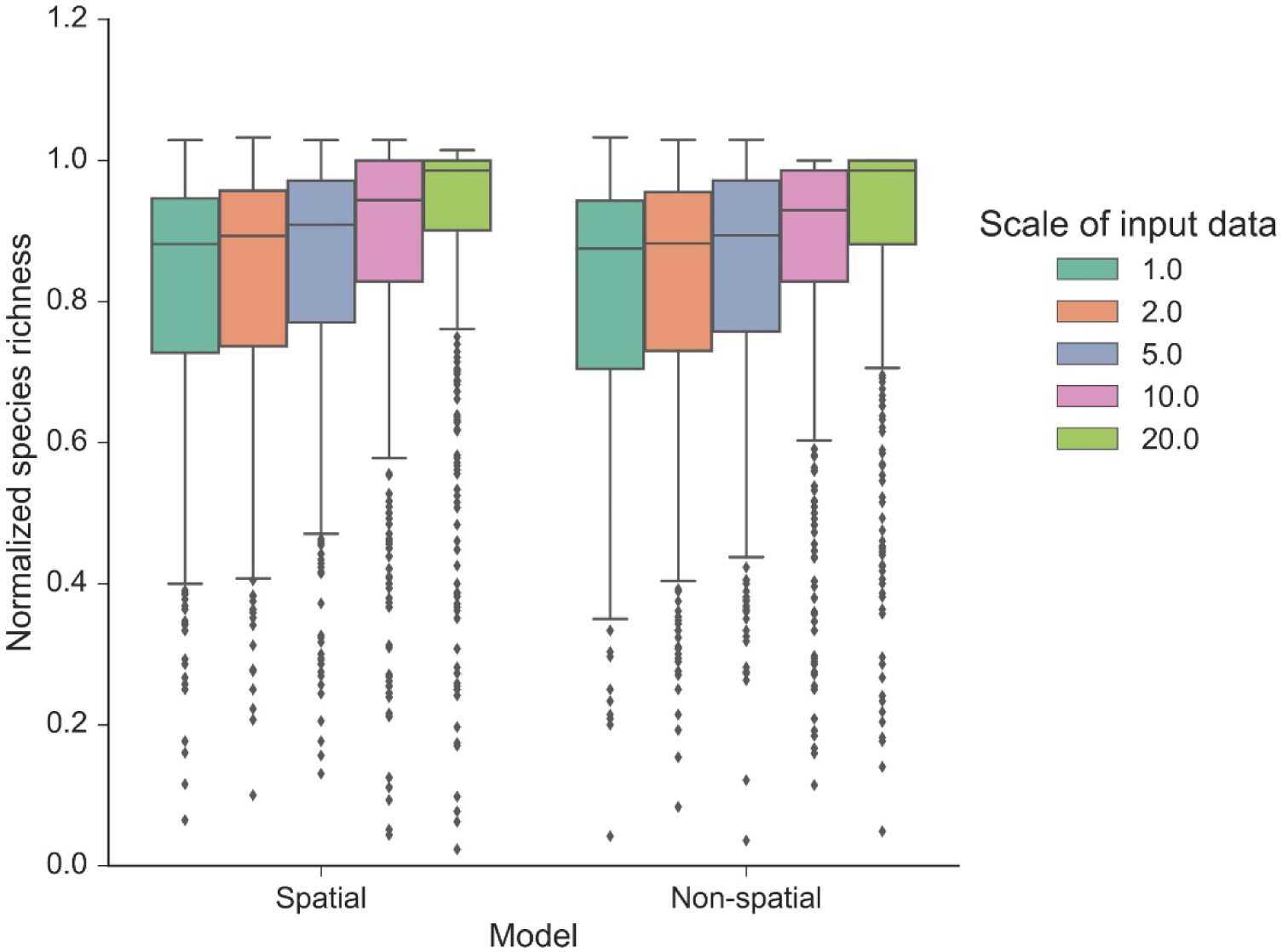
Normalized species richness of the autonomous agent model at various scales. Normalized species richness is the species richness change from time zero compared to the last time step (year 50) of each model run using the sandhills habitat data. Normalized species richness is illustrated between scales of input data and spatial vs. non-spatial dispersal models.

**Fig. 3.**
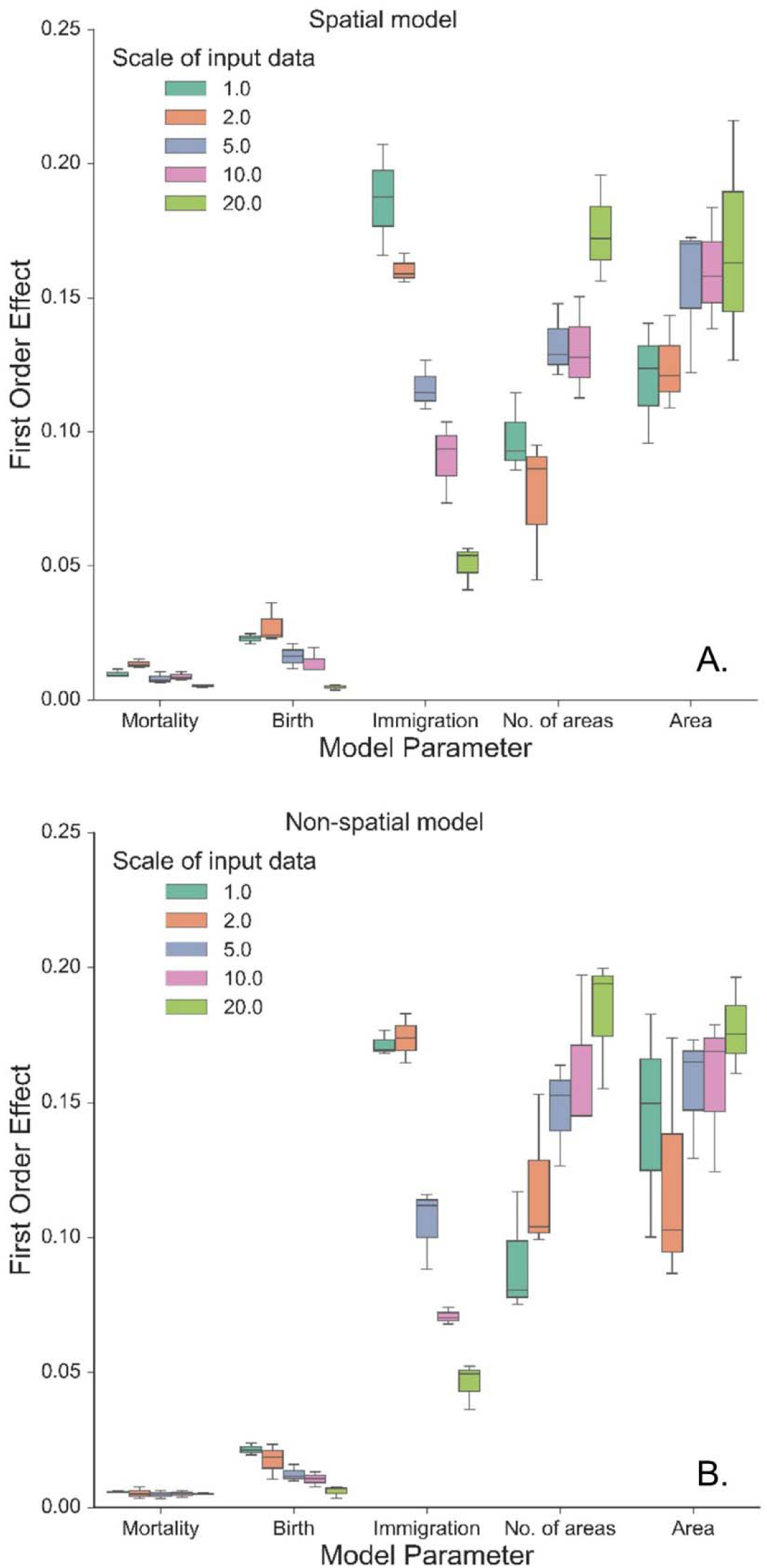
First order effects (%) of the three neutral parameters (mortality, birth, and immigration) and spatial boundary parameters (area and number of areas) on simulated normalized richness using different scales of input data from the sandhills habitat using the spatial model (A.) and the non-spatial model (B.). The legend illustrates the resolution (in dm) at which the data were created from the empirical data, including the original resolution (1.0: 10 cm × 10 cm). Boxplots represent variance across three replicates of the model run using 500 FAST parameter combinations.

The model was sensitive to the scale of the input frequency distributions (**Fig. 2**) and had mixed effects on individual parameter sensitivity (**Fig. 3**). At the finest scales of input distributions (10 cm ×10 cm), immigration rate (compared to birth and mortality) had the largest individual parameter sensitivity on normalized richness (18-20% vs. <3%). Generally, as scale of input distributions coarsened, the neutral parameter sensitivities decreased, particularly for immigration. Effects of scale on mortality rates were minor, with little variability across scales of input data, and low overall individual parameter sensitivity. Scale had significant consequences for the sensitivity of both community size parameters. In general, as the input scale coarsened, the individual parameter sensitivity increased, rather than decreased, as seen with the neutral parameters. The FAST also illustrated some non-linear differences in sensitivity across scales for the community size parameters (e.g. “area” parameter in **Fig. 3B**).

## 4 Discussion

This modeling experiment provided support for neutral processes as a mechanism driving groundcover community dynamics in this high fire frequency ecosystem. This is the first study to examine explicitly fine-scale plant demographics; prior research focused on correlations at the habitat level (Kirkman et al. 2001, Pausas and Ribeiro 2017) and plot level manipulation experiments on species competitive traits (Myers and Harms 2009, Iacona et al. 2012). These findings point to the importance of neutral processes in supporting the high species diversity in longleaf pine ecosystems and they help explain the tight coupling among fire frequency, species richness, and the abatement of competition. This study also supports the central assumption of the UNTB: that niche-based processes do not drive community assembly in these groundcover communities when fire frequency is high.

We simulated a range of groundcover species richness distributions (**Fig. 2**) using simple probability functions, representative of the UNTB. We found that a broad range of estimates for mortality and birth (values from 0.2 to 0.6) can be used to simulate species richness. This suggests that robust estimates are produced without including individual species and their respective traits, and efforts to obtain better estimates may be unnecessary. However, the high sensitivity of dispersal limitation (immigration) illustrates the importance of quantifying accurate immigration rates and of understanding community level dispersal when modeling fine-scale community richness. The importance of immigration, effects on species turnover, the difficulty in its parameterization, and the underlying assumptions were similar to other UNTB simulated systems (Chesson 2000, Girdler and Barrie 2008, Lowe and McPeek 2014, Matthews and Whittaker 2014) and should be explored further.

We found that simulated community stability and rarity were influenced by the community size parameters that affected population size and feedbacks with the neutral parameters (**Fig. 3**). Simulating greater number of areas or larger spatial extents yielded more stable communities and a greater likelihood for rarer species to persist through time when populations are large. The number of areas simulated directly affected the number of local communities, while area simulated affected size of the local community pool. Furthermore, the low effects from different functions of dispersal (spatial vs. non-spatial dispersal, **Figs. 2, 3**) suggests that defining dispersal processes may be trivial when modeling species richness at these fine-scales. In reality, dispersal distance is important for recruitment processes generating and maintaining understory plant diversity. As such, restricting recruitment in the local community using the spatial model can create a modeling environment conducive to realistic dispersal patterns of these small plants (Stamp and Lucas 1990, Kirkman et al. 2004). Here, various dispersal distances and functions can be manipulated within the model and explored across sites. These spatial dispersal processes may be particularly important at coarser scales, where dispersal distance is more of a relevant process, which applies to mosaics of forest management and fire regimes across the landscape.

Population structures, represented here as different input species frequency distributions derived from different scales, were also important (**Fig.s 2, 3**). At coarser scales, the input frequency data were less skewed, where rare species were no longer comparatively rare and simulated richness was more stable than using finer scale data. The effects of scale on each individual parameter was complex (**Fig. 3**). At the finest scales of input distributions, immigration rate had the largest individual sensitivity among the neutral parameters affecting normalized richness. Here, rarer species depended more on immigration from the metacommunity than using coarser scaled inputs. The lower individual sensitivities of birth rate illustrated that rarer species were more dependent on effective recruitment at finer scales than at coarser scales. The overall low sensitivity of mortality illustrated that at even high mortality rates (here, up to 0.6), the community would thrive if the local and metacommunity pool were abundant and available. As such, even if birth rates were low, high mortality values would provide more space – open cells – available for new recruits.

This study illustrated the relative importance of our neutral model parameters and how population structure and spatial boundaries can influence simulated species richness. Our next step is to critically examine the multi-scalar outputs to the empirical data and examine which parameter combinations best predict high species richness. Furthermore, these results do not illustrate the spatial or temporal scales at which neutral processes might break down, specifically where there are changes in fire frequency, land use, overstory structure, fuel characteristics, and soil properties, as well as variability in fire intensity within and among burns. These effects of scale on community structure and rarity are however consistent with both neutral theory and island biogeography predictions (Volkov et al. 2003, MacArthur and Wilson 2016) and provide an example of patterns of diversity maintained by both processes. Furthermore, the fine-scale quantification of frequency distributions for these groundcover species represents the spatial scale (100 cm^2^) at which community dynamics occur, which was important for examining model simulations and parameter sensitivity.

An alternative approach of characterizing over 100 plant species by their life history traits and incorporating them into a niche-based model (Peterson 2006) would be significantly more complex than this simple modeling approach. This would likely provide little to no improvement in predicting species richness patterns under high fire frequencies, particularly if stochastic processes are driving community dynamics. Utilizing species traits would be useful however, when quantifying the extent to which niche-based processes determine community dynamics when fire frequency is altered and in examining the impact of variation of fire intensity and resulting burn severity within and among burns (O’Brien et al. 2016a, b). A deeper understanding of the impact of variable fire regimes would require combining a neutral model with a competitive exclusion model operating at the functional group level. This strategy has been explored when examining the impact of stochastic versus deterministic processes across ecosystems (defined as the Continuum Hypothesis, Gravel et al. 2006). For this more inclusive framework, deterministic and neutral assumptions form the ends of a gradient, defined in this case by fire frequency or spatial scale. However, we argue that the Continuum Hypothesis plays out within a single landscape driven by the dynamics of fire. When fire is removed, the transition from neutral to deterministic community dynamics can also vary across landscapes, with competitive exclusion occurring more quickly in higher productivity sites than on less productive xeric sites (Kirkman et al. 2001).

Frequently burned ecosystems represent a reservoir of biodiversity worldwide, and it is critical to understand the processes by which fire maintains diversity. This will aid in supporting critical elements of managed fire regimes, setting realistic restoration targets, and examining monitored changes in plant communities (Hiers et al. 2012). We found that simple neutral-based simulations were useful for projecting species richness at multiple scales, although further work is needed to determine optimal parameterization and to test against empirical data. These results are globally relevant because similar woodland and grassland ecosystems that are structured by the obligate process of frequent fire occur worldwide, and all are threatened by alterations to their fire regime. Examples include tallgrass prairies of North America (Leach and Givnish 1996), African savannas (Bond et al. 2005), Australian eucalypt woodlands (Bowman 2000), and cerrado savannas of Brazil (Mistry 1998). These fire-maintained ecosystems may be governed by neutral processes generated by fine-scale fire mortality and fire driven competitive abatement to maintain diversity in their groundcover communities.

## 5 Acknowledgments

We acknowledge the Eglin Air Force Base Natural Resource Management Branch and the Air Force Wildland Fire Center, Niceville, FL for their support and technical assistance, particularly B. Williams. We acknowledge the USDA Forest Service, Southern Research Station and the Center for Forest Disturbance Science, Athens, GA for their support.

## 6 Funding

This research was funded by the US Department of Defense Strategic Environmental Research and Development Program (#RC-2243) and the National Science Foundation (DEB-1442103). This work was also supported by the US Department of Agriculture Forest Service National Fire Plan.

## 7 Author Contributions Statement

LL, LD, JO, SP, AH, BH, LR, JD, SG, and KH contributed to manuscript writing and advised in the development and interpretation of the autonomous agent model, sampling design, and analyses. LL was the lead in model development, FAST analysis, and manuscript writing, with significant contributions by LD and JO. All authors contributed to idea development and implementation. Other contributions include, but are not limited to, field plant monitoring by SP; sensitivity analysis by SG; and prescribed burning by BH and KH.

