## Supplementary Material for "Simulating groundcover community assembly in a frequently burned ecosystem using a simple neutral model"

**Table S1.** Plant species list for each habitat monitored at EAFB for this study, ordered alphabetically.

| Flatwoods | Sandhills |
| --- | --- |
| *Ageratina aromatica* | *Andropogon virginicus* |
| *Andropogon virginicus* | *Aristida mohrii* |
| *Aristida mohrii* | *Aristida purpurascens* |
| *Aristida purpurAscens* | *Balduina angustifolia* |
| *Aristida stricta* | *Bulbostylis ciliatifolia* |
| *Arundinaria gigantea* | *Chrysopsis gossypina* |
| *Bigelowia nudata* | *Cnidoscolus stimulosus* |
| *Bulbostylis ciliatifolia* | *Commelina erecta* |
| *Carphephorus odoratissimus* | *Crataegus michauxii* |
| *Chamaechrista fasciculata* | *Croton argyranthemus* |
| *Chamaechrista nictitans* | *Cyperus filiculmis* |
| *Chrysopsis gossypina* | *Cyperus retrorsus* |
| *Cnidoscolus stimulosus* | *Dichanthelium aciculare* |
| *Commelina erecta* | *Dichanthelium acuminatum* |
| *Croton argyranthemus* | *Dichanthelium ovale* |
| *Crotalaria purshii* | *Dichanthelium portoricens* |
| *Ctenium aromaticum* | *Dichanthelium sphaerocarpon* |
| *Cyperus filiculmis* | *Dichanthelium strigosum* |
| *Dichanthelium aciculare* | *Diodia virginiana* |
| *Dichanthelium ovale* | *Eriogonum tomentosum* |
| *Dichanthelium portoricens* | *Euphorbia floridana* |
| *Dichanthelium ravenelii* | *Euthamia caroliniana* |
| *Dichanthelium sphaerocarpon* | *Galactia erecta* |
| *Dichanthelium strigosum* | *Galactia regularis* |
| *Diodia teres* | *Gaylussacia dumosa* |
| *Elephantopus elatus* | *Helianthemum carolinianum* |
| *Eriogonum tomentosum* | *Hieracium gronovii* |
| *Euphorbia discoidalis* | *Houstonia procumbens* |
| *Euphorbia floridana* | *Hypericum gentianoides* |
| *Galactia regularis* | *Hypoxis juncea* |
| *Gaylussacia dumosa* | *Ionactis linariifolia* |
| *Gaylussacia frondosa* | *Lechea sessiliflora* |
| *Gelsemium sempervirens* | *Liatris graminifolia* |
| *Helianthus radula* | *Liatris secunda* |
| *Hypericum gentianoides* | *Liatris tenuifolia* |
| Flatwoods (continued) | Sandhills (continued) |
| *Hypoxis juncea* | *Licania michauxii* |
| *Ilex cassine* | *Lupinus diffusus* |
| *Ilex decidua* | *Mimosa quadrivalvis* |
| *Ilex glabra* | *Opuntia humifusa* |
| *Ilex vomitoria* | *Paronychia patula* |
| *Ionactis linariifolia* | *Paspalum praecox* |
| *Juncus dichotomus* | *Physalis arenicola* |
| *Kalmia hirsuta* | *Pinus clausa* |
| *Lachnocaulon beyrichianum* | *Pinus palustris* |
| *Lechea sessiliflora* | *Pityopsis aspera* |
| *Lespedeza repens* | *Pityopsis graminifolia* |
| *Liatris elegans* | *Polygonella gracilis* |
| *Liatris graminifolia* | *Polygala polygama* |
| *Liatris tenuifolia* | *Polygala polygama* |
| *Licania michauxii* | *Pteridium aquilinum* |
| *Mimosa quadrivalvis* | *Quercus geminata* |
| *Myrica cerifera* | *Quercus incana* |
| *Opuntia humifusa* | *Quercus laevis* |
| *Paspalum praecox* | *Rhexia mariana* |
| *Pinus elliottii* | *Rhus copallinum* |
| *Pinus palustris* | *Rhynchosia cytisoides* |
| *Pityopsis aspera* | *Rhynchospora grayi* |
| *Pityopsis graminifolia* | *Rubus cuneifolius* |
| *Pityopsis oligantha* | *Schizachyrium scoparium* |
| *Polygonella gracilis* | *Schizachyrium tenerum* |
| *Polygala nana* | *Scleria ciliata* |
| *Polygala polygama* | *Scleria triglomerata* |
| *Polypremum procumbens* | *Serenoa repens* |
| *Quercus incana* | *Smilax auriculata* |
| *Quercus minima* | *Smilax bona-nox* |
| *Rhexia alifanus* | *Solidago odora* |
| *Rhexia lutea* | *Sorghastrum secundum* |
| *Rhynchospora baldwinii* | *Sporobolus junceus* |
| *Rhynchosia cytisoides* | *Stylosanthes biflora* |
| *Rhynchospora grayi* | *Stylisma patens* |
| *Rubus cuneifolius* | *Tephrosia chrysophylla* |
| *Schizachyrium scoparium* | *Tephrosia virginiana* |
| *Schizachyrium tenerum* | *Tradescantia hirsutiflora* |
| *Scleria ciliata* | *Tragia smallii* |
| *Scleria triglomerata* | *Tragia urens* |
| Flatwoods (continued) | Sandhills (continued) |
| *Serenoa repens* | *Vaccinium arboreum* |
| *Seymeria cassioides* | *Vaccinium darrowii* |
| *Smilax auriculata* |  |
| *Smilax laurifolia* |  |
| *Smilax smallii* |  |
| *Solidago odora* |  |
| *Sorghastrum secundum* |  |
| *Stylosanthes biflora* |  |
| *Stylisma patens* |  |
| *Symphyotrichum dumosum* |  |
| *Tephrosia chrysophylla* |  |
| *Tephrosia florida* |  |
| *Tephrosia hispidula* |  |
| *Tephrosia virginiana* |  |
| *Tragia smallii* |  |
| *Tragia urens* |  |
| *Vaccinium corymbosum* |  |
| *Vaccinium darrowii* |  |
| *Vernonia angustifolia* |  |
| *Viola palmata* |  |
| *Xyris brevifolia* |  |
| *Xyris caroliniana* |  |
| *Xyris elliottii* |  |

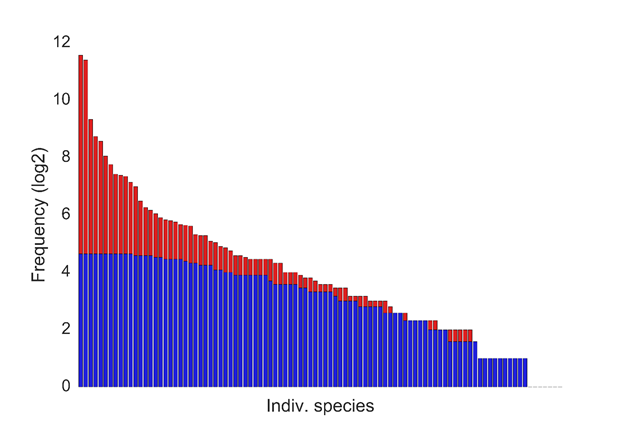

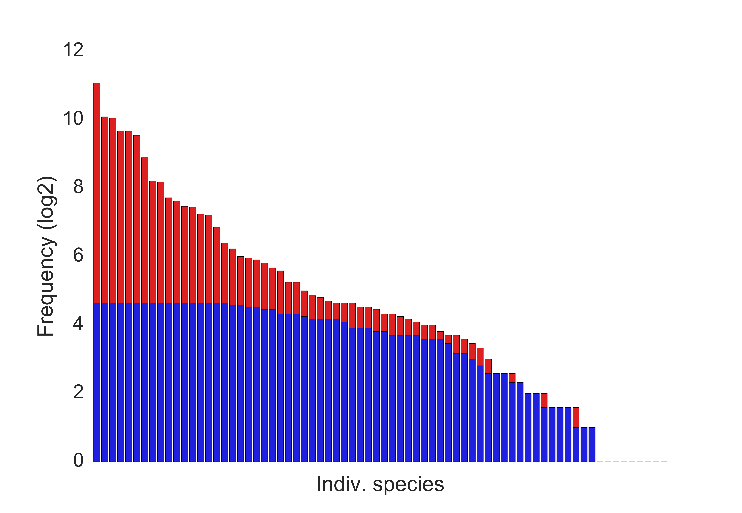

**Figure S1.** Example input frequency distribution in log_2_ scale based on scaling the input data by coarsening the 10 cm x 10 cm data. Red: original resolution (10 cm x 10 cm). Blue: scaled to 200 cm x 200 cm. Flatwoods distribution on the left; Sandhills on the right.

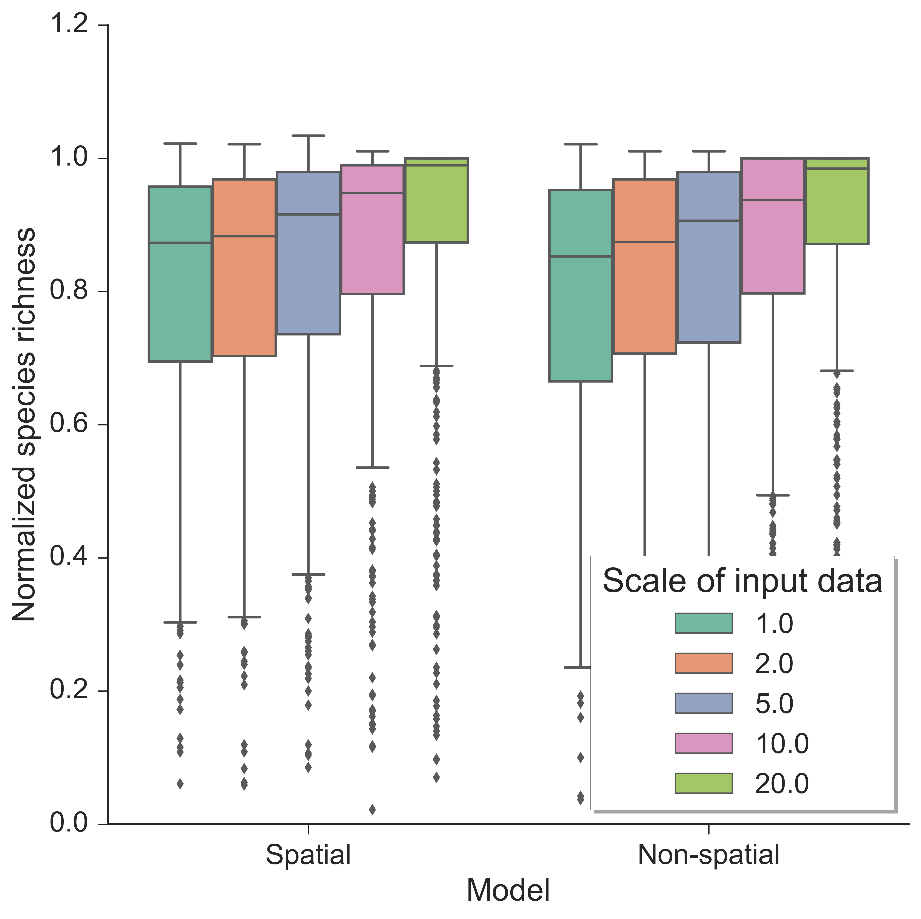

**Figure S2.** Normalized species richness of the autonomous agent model at various scales. Normalized species richness is the species richness change from time zero compared to the last time step (year 50) of each model run using the flatwoods habitat data. Normalized species richness is illustrated between scales of input data and spatial vs. non-spatial dispersal models.

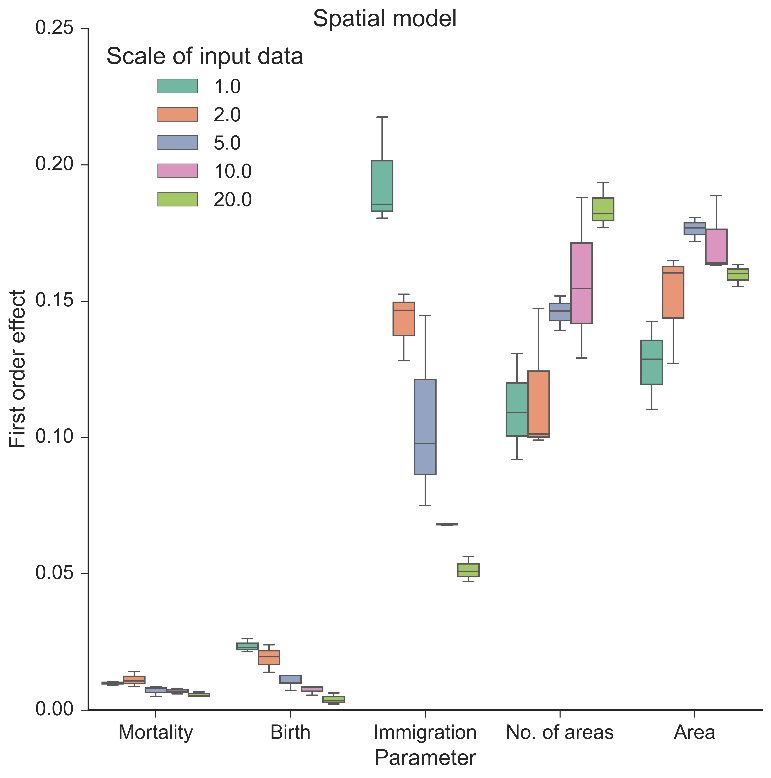
 **A.**

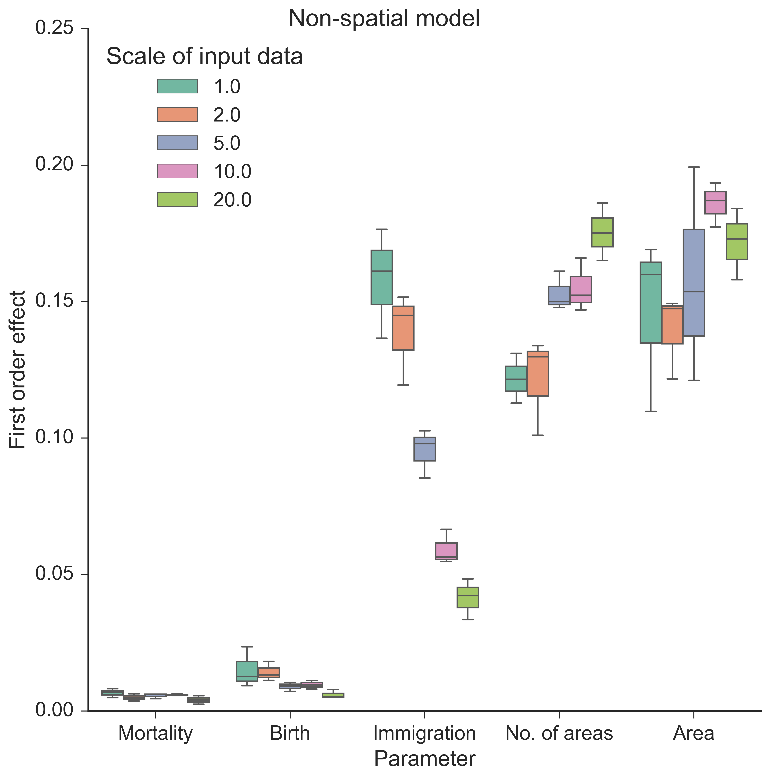
 **B.**

**Figure S3.** First order (%) effect of the three neutral parameters (mortality, birth, and immigration) and spatial boundary parameters (area and number of areas) on simulated normalized richness using different scales of input data in the flatwoods habitat using the spatial model (A.) and non-spatial model (B.). The legend illustrates the resolution (in dm) in which the data was created from the empirical data, including the original resolution of the data (1.0: 10 cm x 10 cm). Boxplots represent variance across three replicates of the model run using 500 FAST parameter combinations.
